# A chromosome-level genome assembly for the Eastern Fence Lizard (*Sceloporus undulatus*), a reptile model for physiological and evolutionary ecology

**DOI:** 10.1101/2020.06.06.138248

**Authors:** Aundrea K. Westfall, Rory S. Telemeco, Mariana B. Grizante, Damien S. Waits, Amanda D. Clark, Dasia Y. Simpson, Randy L. Klabacka, Alexis P. Sullivan, George H. Perry, Michael W. Sears, Christian L. Cox, Robert M. Cox, Matthew E. Gifford, Henry B. John-Alder, Tracy Langkilde, Michael J. Angilletta, Adam D. Leaché, Marc Tollis, Kenro Kusumi, Tonia S. Schwartz

## Abstract

High-quality genomic resources facilitate population-level and species-level comparisons to answer questions about behavioral ecology, morphological and physiological adaptations, as well as the evolution of genomic architecture. Squamate reptiles (lizards and snakes) are particularly diverse in characteristics that have intrigued evolutionary biologists, but high-quality genomic resources for squamates are relatively sparse. Lizards in the genus *Sceloporus* have a long history as important ecological, evolutionary, and physiological models, making them a valuable target for the development of genomic resources. We present a high-quality chromosome-level reference genome assembly, SceUnd1.0, (utilizing 10X Genomics Chromium, HiC, and PacBio data) and tissue/developmental stage transcriptomes for the Eastern Fence Lizard, *Sceloporus undulatus*. We performed synteny analysis with other available squamate chromosome-level assemblies to identify broad patterns of chromosome evolution including the fusion of micro- and macrochromosomes in *S. undulatus*. Using this new *S. undulatus* genome assembly we conducted reference-based assemblies for 34 other *Sceloporus* species to improve draft nuclear genomes assemblies from 1% coverage to 43% coverage on average. Across these species, typically >90% of reads mapped for species within 20 million years divergence from *S. undulatus*, this dropped to 75% reads mapped for species at 35 million years divergence. Finally we use RNAseq and whole genome resequencing data to compare the three assemblies as references, each representing an increased level of sequencing, cost and assembly efforts: Supernova Assembly with data from10X Genomics Chromium library; HiRise Assembly that added data from HiC library; and PBJelly Assembly that added data from PacBio sequencing. We found that the Supernova Assembly contained the full genome and was a suitable reference for RNAseq, but the chromosome-level scaffolds provided by the addition of the HiC data allowed the reference to be used for other whole genome analysis, including synteny and whole genome association mapping analyses. The addition of PacBio data provided negligible gains. Overall, these new genomic resources provide valuable tools for advanced molecular analysis of an organism that has become a model in physiology and evolutionary ecology.

## Context

Genomic resources, including high-quality reference genomes and transcriptomes, facilitate comparisons across populations and species to address questions ranging from broad-scale chromosome evolution to the genetic basis of key adaptations. Squamate reptiles, the group encompassing lizards and snakes, have served as important models in ecological and evolutionary physiology due to their extensive metabolic plasticity [1]; diverse reproductive modes including obligate and facultative parthenogenesis [2]; repeated evolution of placental-like structures [2, 3]; shifts among sex determining systems, with XY, ZW, and temperature-dependent systems represented often in closely related lizards species [4, 5]; loss of limbs and elongated body forms [6]; and the ability to regenerate tissue [7, 8].

Despite having evolved greater phylogenetic diversity than mammals and birds, two major vertebrate groups with extensive genome sampling, genomic resources for squamates remain scarce and assemblies at the chromosome-level are even more rare [7, 9-13]. While squamates are known to have a level of karyotypic variability similar to that of mammals [14], the absence of high-quality genome assemblies has led to their exclusion from many chromosome-level comparative genome analyses. In comparative studies, non-mammalian amniotes are often represented only by the chicken, which is divergent from squamate reptiles by almost 280 million years [15], or the green anole (*Anolis carolinensis*), whose genome is only 60% assembled into chromosomes and is lacking assembled microchromosomes [14, 16]. However, recent analyses have identified key differences that distinguish the evolution of squamate genomes from patterns found in mammals and birds [17], underscoring the need for additional high-quality genome assemblies for lizards and snakes. The development of additional squamate genomes within and across lineages will facilitate investigations of the genetic basis for many behavioral, morphological, and physiological adaptations in comparisons of organisms from the population up to higher-order taxonomic ranks.

Our goal was to develop a high-quality genomic and transcriptomic resources for the spiny lizards (*Sceloporus*) to further our ability to address fundamental ecological and evolutionary questions within this taxon, across reptiles and across vertebrates. The genus *Sceloporus* includes approximately 100 species extending throughout Central America, Mexico, and the United States [18]. Researchers have used *Sceloporus* for decades as a model system in the study of physiology [19, 20], ecology [21, 22], reproductive ecology [23-25], life history [26-28], and evolution [25, 29-31]. The long history of research on *Sceloporus* species, applicability across multiple fields of biology, and the extensive diversity of the genus makes this an ideal group to target for genomic resource development.

We focus on the Eastern fence lizard, *Sceloporus undulatus*, which is distributed in forested habitats east of the Mississippi River [32]. Recently, *S. undulatus* has been the focus of studies on the development of sexual size dimorphism [33, 34], as well as experiments testing the effects of invasive species [35-37] and climate change [22, 38-40] on survival and reproduction as a model to understand better the broader consequences of increasing anthropogenic disturbance. The development of genomic resources for S. undulates, particularly a high-quality genome assembly, will support its role as a model species for evolutionary and ecological physiology, and will have immediate benefits for a broad range of comparative studies in physiology, ecology, and evolution.

To this end, we developed a high-quality chromosome-level reference genome assembly and transcriptomes from multiple tissues for the *S. undulatus*. We apply this genome reference to datasets on three scales: (1) to address how assembly quality influences mapping in RNAseq and low coverage whole-genome sequence data; (2) to improve upon the genomic resources for the *Sceloporus* genus by creating reference-based assembly of draft genomes for 34 other *Sceloporus* species; and (3) to draw broad comparisons in chromosome structure and conservation with other recently published squamate chromosome-level genomes through large-scale synteny analysis.

## Methods and Analyses

### Sequencing and assembly of the *Sceloporus undulatus* genome

Genome sequence data were generated from two male individuals collected at Solon Dixon Forestry Education Center, in Andalusia, Alabama (31°09’49”N, 86°42’10”W). The animals were euthanized and tissues were dissected, snap-frozen in liquid nitrogen, and stored at – 80°C. Procedures were approved by the Pennsylvania State University Institutional Animal Care and Use Committee (Protocol# 44595-1).

We developed three *S. undulatus* genome assemblies using increasingly more data with correspondingly greater cost: (1) a SuperNova assembly containing data from 10X Genomics Chromium, (2) a HiRise assembly containing the 10X Genomics data with the addition of Hi-C data, and (3) a PBJelly Assembly containing the 10X Genomics data and Hi-C data, and the addition of PacBio data. These assemblies are provided as supplemental files and their summary statistics are provided in Table 1.

**Table 1.**
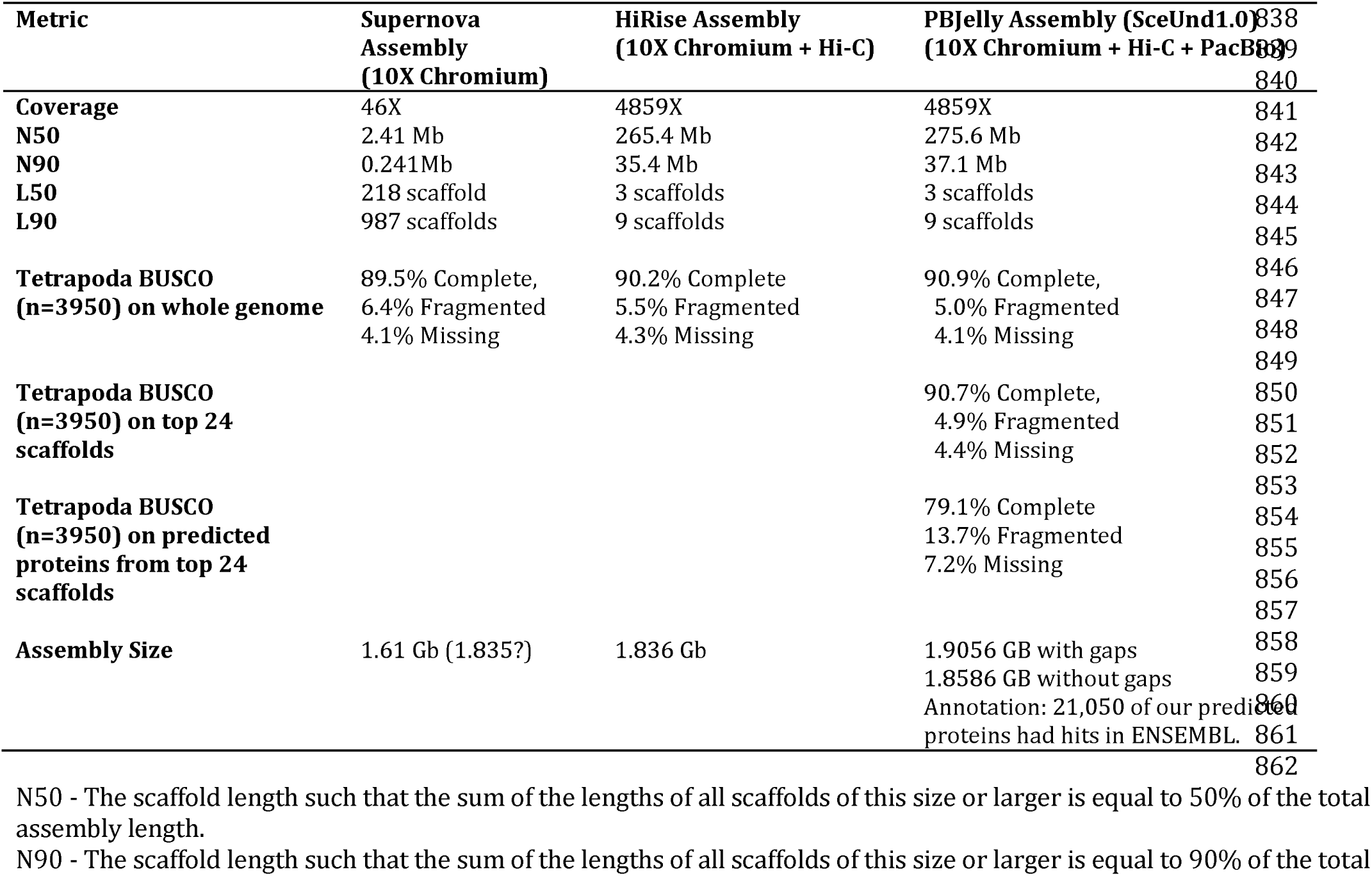
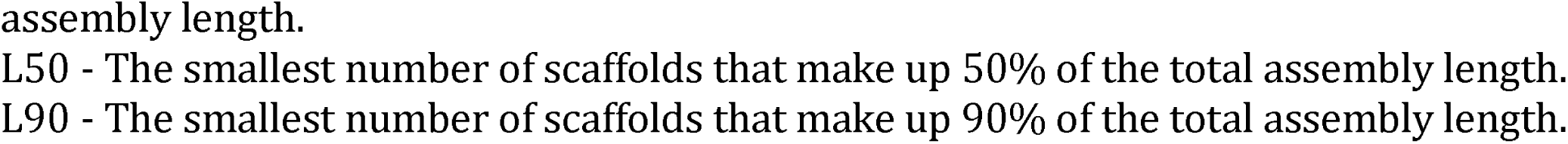
Summary statistics across genome assemblies.

In the fall of 2016, we sequenced DNA from snap-frozen brain tissue of a single juvenile male *S. undulatus* using 10X Genomics Chromium Genome Solution Library Preparation with SuperNova Assembly [41] through HudsonAlpha. The library was sequenced on one lane of Illumina HiSeqX resulting in 774 million 150 bp paired-end reads that were assembled using the SuperNova pipeline. We refer to this assembly as the SuperNova Assembly.

In the fall of 2017, we sequenced a second male (Figure 1) from the same population using a Hi-C library with Illumina sequencing through Dovetail Genomics prepared from blood, liver, and muscle tissue. The remains from the first individual that was used for the SuperNova Assembly were insufficient for the Hi-C library preparation, which required 100 mg of tissue. Dovetail Genomics developed two Hi-C libraries that were sequenced on an Illumina HiSeqX to produce 293 million and 289 million (total 582 million) 150 bp PE reads. The data from both the Hi-C and the 10X Genomics were used for assembly in the HiRise software pipeline at Dovetail Genomics. We refer to this as the HiRise Assembly.

**Figure 1.**
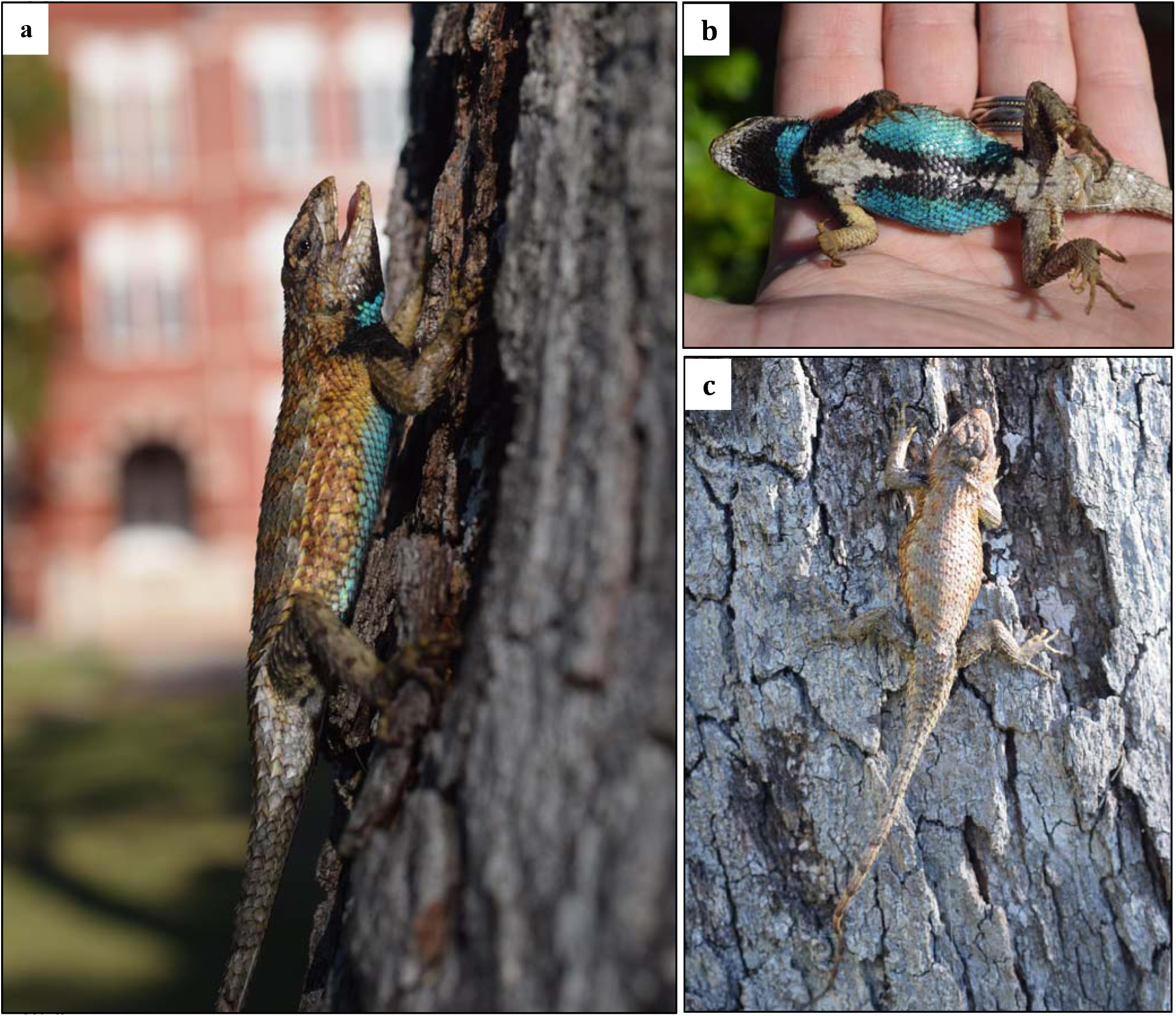
Adult male *Sceloporus undulatus* (Eastern Fence Lizard) from Andalusia, Alabama, pictured outside of Sanford Hall at Auburn University, (a) profile, (b) ventral, (c) dorsal view. This specimen was used for genome sequencing at DoveTail Genomics. Photo credits to R. Telemeco.

Finally, also in fall of 2017, DNA extracted from the same adult male individual was used by Dovetail Genomics to generate 1,415,213 PacBio reads with a mean size of 12,418.8 bp (range 50-82,539 bp). These PacBio data were used for gap-filling to further improve the lengths of the scaffolds of the HiRise Assembly using the program PBJelly [42]. We refer to this final assembly containing all three types of sequencing data as the PBJelly Assembly and the SceUnd1.0 reference genome assembly.

For a visual comparison among the three assemblies and to other squamate genomes, we graphed the genome contiguity for these three assemblies with other squamate reptile genomes, building on the graph by Roscito et al. [42]. The Eastern fence lizard, *S. undulatus*, SuperNova Assembly (containing only the 10X Genomics data) is as contiguous as the bearded dragon genome assembly (Figure 2a). The addition of the HiRise data brought a large increase in continuity. The HiRise and PBJelly *S. undulatus* Assemblies and are nearly indistinguishable from each other and are among the most contiguous squamate genome assemblies to date (Figure 2a).

**Figure 2.**
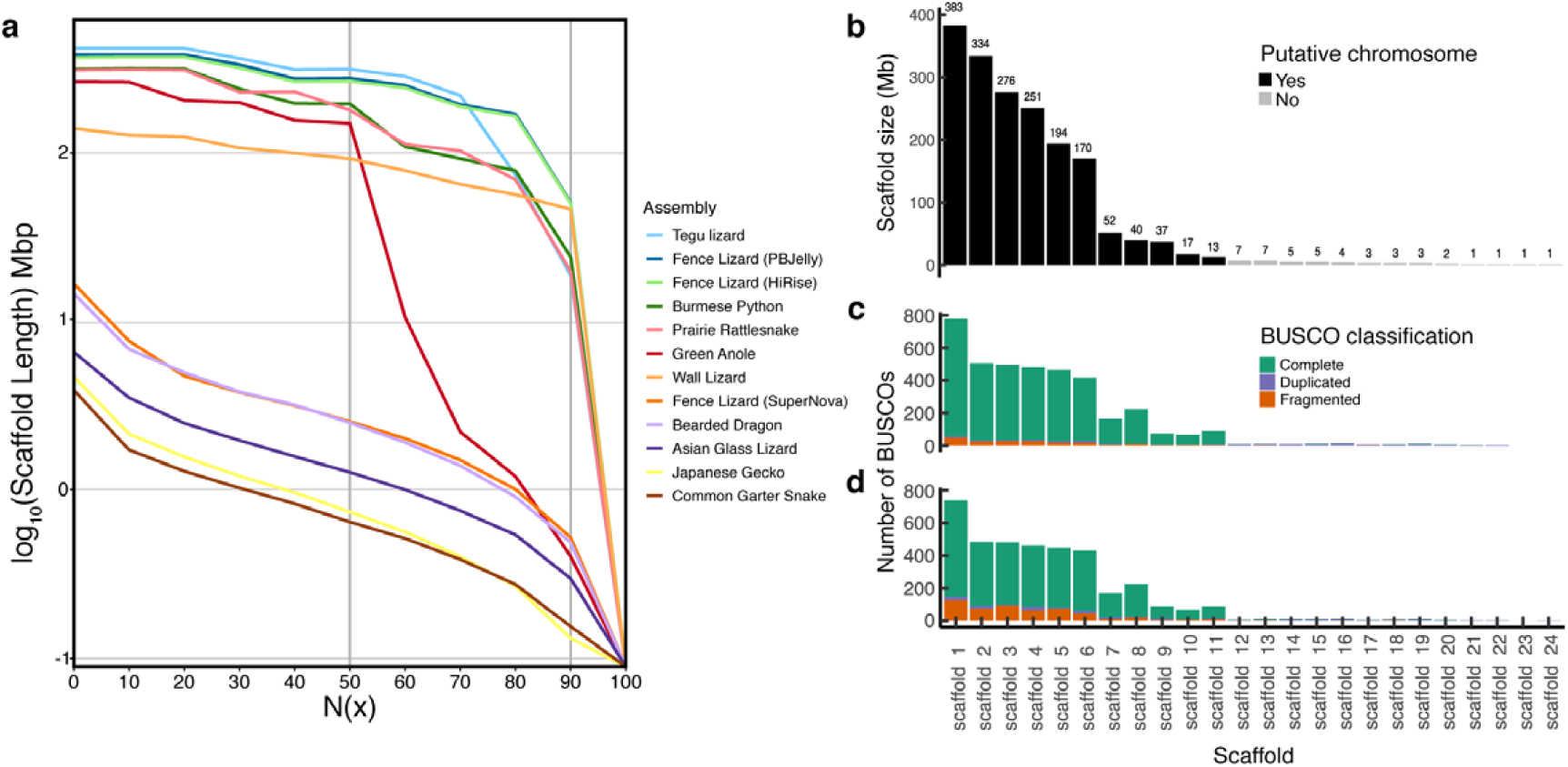
An evaluation of *S. undulatus* genome assembly quality. (a) Comparison of the contiguity of the three *S. undulatus* genome assemblies (Fence Lizard) relative to other squamates genome assemblies based on the log 10 of the scaffold length. The X axis is the N(x) with the N50 and the N90 emphasized with a vertical line, representing the scaffold size that contains 50 or 90 percent of the data. The legend lists the assemblies in the order of the lines from most contiguous (top) to least contiguous (bottom). Note the Fence Lizard PBJelly (dark blue, SceUnd1.0) and Fence Lizard HiRise (green) assemblies are the second and third from the top and are nearly indistinguishable. (b-d) Scaffold size distribution of SceUnd1.0 and the number of BUSCO genes that mapped to each scaffold. The length of the first 24 scaffolds, where the first 11 scaffolds likely represent the haploid N=11 chromosomes (6 macrochromosomes and 5 microchromosomes). The numbers above each bar represent scaffold length to the nearest Mb. The number of BUSCO genes that mapped to each scaffold based on (c) the genome assembly, and (d) the predicted proteins from the annotation. The 11 large scaffolds inferred to correspond to chromosomes have many unique and complete BUSCO genes (green), whereas the smaller contigs have many duplicated BUSCOs (purple) suggesting they are the result of reads not mapping correctly to the chromosomes.

The SceUnd1.0 assembly contains 45,024 scaffolds (>850 bp, without gaps) containing 1.9 Gb of sequence, with N50 of 275 Mb. Importantly, 92.6% (1.765 Gb) of the assembled sequence is contained within the first 11 scaffolds. Chromosomal studies have determined that the *S. undulatus* karyotype is 2N = 22 with a haploid genome of N = 11 (six macrochromosomes + five microchromosomes; 6M + 5m) [31, 43]. Sorting the top 11 scaffolds by size (Figure 2b) suggests that scaffolds 1-6 are the macrochromosomes (170-383 Mb in size) and scaffolds 7-11 are the five microchromosomes (13-52 Mb in size) (Figure 2b). These results suggest that the first 11 scaffolds represent the 11 chromosomes, although the assembly also produces 45,000 tiny scaffolds between 0.85KB – 7MB that may still contain relevant chromosomal segments that could not be assembled.

To assess the completeness of the three genome assemblies, we utilized the BUSCO (Benchmarking Universal Single-Copy Orthologues) Tetrapoda dataset (3950 genes) [44, 45]. For all three assemblies we found over 89% of BUSCO genes complete (Table 1) with only minor differences in BUSCO genes between the SuperNova, HiRise, and PBJelly Assemblies (89.5%, 90.2%, 90.9% complete). This suggests that the initial SuperNova Assembly captured nearly all of the genomic content despite having considerably shorter scaffolds (Table 1). The small increase in success with the more contiguous assemblies appears to be the result of a reduction in fragmented BUSCO genes with increasing data. In the SuperNova Assembly 6.4% of BUSCO genes were present as fragments whereas only 5.5% and 5.0% are present as fragments in the HiRise and PBJelly Assemblies, respectively, thus explaining the 1.4% difference in complete BUSCO genes present. Interestingly, there was a 0.2% (i.e., 8 genes) increase in missing BUSCO genes from the SuperNova to the HiRise Assembly. In the PBJelly Assembly (SceUnd1.0), the BUSCO genes are almost all found on the largest 11 scaffolds (Figure 2c), as we would predict if those scaffolds correspond to chromosomes. Most of the BUSCO genes on the smaller scaffolds were duplicated. Even so, there are a small number of complete and fragmented BUSCO genes present on a handful of the tiny scaffolds (Figure 2c), suggesting that these scaffolds contain pieces of the chromosomes that were not properly assembled.

### De novo assembly and annotation of the *Sceloporus undulatus* transcriptome

Samples used for the *de novo* transcriptome were obtained from three gravid females of *Sceloporus undulatus* collected in Edgefield County, South Carolina (33.7°N, 82.0°W) and transported to Arizona State University. These animals were maintained under conditions described in previous publications [46, 47], which were approved by the Institutional Animal Care and Use Committee (Protocol #14-1338R) at Arizona State University. Approximately two days after laying eggs, each lizard was euthanized by injecting sodium pentobarbital into the coelomic cavity. Whole brain and skeletal muscle samples were removed and placed in RNA-lysis buffer (mirVana miRNA Isolation Kit, Ambion) and flash-frozen. Additionally, three early-stage embryos from each clutch were dissected, pooled together, homogenized in RNA-lysis buffer, and also flash frozen.

Total RNA was isolated from the embryo and three tissue samples from each adult female (whole brain, skeletal muscle) using the mirVana miRNA Isolation Kit (Ambion) total RNA protocol. Samples were checked for quality on a 2100 Bioanalyzer (Agilent). One sample from each tissue was selected for RNAseq based on the highest RNA Integrity Number (RIN), with a minimum cutoff of 8.0. For each selected sample, 3 μg of total RNA was sent to the University of Arizona Genetics Core (Tucson, AZ) for library preparation with TruSeq v3 chemistry for a standard insert size. RNA samples were multiplexed and sequenced using an Illumina HiSeq 2000 to generate 100-bp paired-end reads. Publicly available raw Illumina RNAseq reads from *S. undulatus* liver (juvenile male) were also added to our dataset [48, 49]. After removing adapters, raw reads from the four tissues were evaluated using FastQC (https://github.com/s-andrews/FastQC) and trimmed using Trimmomatic v-0.32 [50], filtering for quality score (≥Q20) and using HEADCROP:9 to minimize nucleotide bias. This procedure yielded 179,374,469 quality-filtered reads. Table 2 summarizes read-pair counts from whole brain, skeletal muscle, whole embryos, and liver.

**Table 2.** *Sceloporus undulatus de novo* transcriptome assembly statistics. The four tissues are comprised of 3 tissues first reported in this study (brain, skeletal, and embryos) from gravid females collected in Edgefield County, SC), plus liver tissue as previously reported by McGaugh et al. 2015.

| **Assembly** | **1 tissue [23]** | **3 tissues** | **4 tissues** |
| --- | --- | --- | --- |
| Total of Trinity transcripts | 158,323 | 492,249 | 547,370 |
| Total of Trinity 'genes' | 138,031 | 422,687 | 467,658 |
| GC% | 43.81 | 42.85 | 42.76 |
| Contig N50 | 1,720 | 1,648 | 1,438 |
| Contig E90N50 | 2,254 | 2,640 | 2,550 |
| Average contig length (bp) | 833.0 | 822.4 | 781.5 |
| Transcripts with the longest ORFs | 86,630<br>(54.7%) | 212,172<br>(43.1%) | 217,756<br>(39.8%) |

All trimmed reads were pooled and assembled *de novo* using Trinity v-2.2.0 with default k-mer size of 25 [51, 52]. From the final transcriptome, a subset of contigs containing the longest open reading frames (ORFs), representing 123,323 transcripts, was extracted from the *de novo* transcriptome assembly using TransDecoder v-3.0.0 (http://transdecoder.github.io) with homology searches against the databases UniProtKB/SwissProt [53] and PFAM [54]. The transcriptome was annotated using Trinotate v-3.0 (http://trinotate.github.io), which involved searching against multiple databases (as UniProtKB/SwissProt, PFAM, signalP, GO) to identify sequence homology and protein domains, as well as to predict signaling peptides. This pooled Tissue-Embryo Transcriptome and annotation are provided as supplemental files.

The most comprehensive transcriptome, obtained using reads from four tissues, consists of 547,370 contigs with an average length of 781.5 nucleotides (Table 2) — shorter than other assemblies because of the range of contig sizes that varied among datasets (1, 3 and 4 tissues; Table S1, Fig. S1). The N50 of the most highly expressed transcripts that represent 90% of the total normalized expression data (E90N50) was lowest in the assembly based on one tissue (Table 2).

To validate the *de novo* transcriptome data, trimmed reads from the 4 tissues used for RNA sequencing (brain, skeletal muscle, liver and whole embryos) were aligned back to the Trinity assembled contigs using Bowtie2 v2.2.6 [55]. From the 176,086,787 reads that aligned, 97% represented proper pairs (Table S2), indicating good read representation in the *de novo* transcriptome assembly. To assess quality and completeness of the assemblies, we first compared the *de novo* assembled transcripts with the BUSCO Tetrapoda dataset, with BLAST+ v2.2.31 [56] and HMMER v3.1b2 [57] as dependencies. This procedure revealed that the *de novo* transcriptome assembly captured 97.1% of the expected orthologues (sum of completed and fragmented), a result comparable to the 97.8% obtained for the green anole transcriptome using 14 tissues [58] (Table 3). Next, nucleotide sequences of *de novo* assembled transcripts with the longest ORFs were compared to the protein set of *Anolis carolinensis* (AnoCar2.0, Ensembl) using BLASTX (evalue=1e-20, max_target_seqs=1). This comparison showed that 11,223 transcripts of *S. undulatus* have nearly full-length (>80%) alignment coverage with *A. carolinensis* proteins (Table S3). Predicted proteins of *S. undulatus* were also used to identify 13,422 one-to-one orthologs with proteins of *A. carolinensis* through reciprocal BLAST (evalue=1e-6, max_target_seqs=1). Table 4 summarizes the *de novo* transcriptome annotation results.

**Table 3.** BUSCO results for transcriptomes of lizard species. For *S. undulatus*, the 4 tissues are the 3 tissues (brain, skeletal muscle and embryos) with the addition of 1 tissue (liver) from McGaugh et al. 2015. For *A. carolinensis*, see Eckalbar et al. 2013 for the complete list of tissues used.

|  | *Sceloporus undulatus* |  |  | *Anolis carolinensis* |
| --- | --- | --- | --- | --- |
|  | 1 tissue | 3 tissues | 4 tissues | 14 tissues |
| Complete genes | 72.5% | 91.7% | 92.3% | 96.7% |
| Duplicated genes | 25% | 43.8% | 43.9% | 37.9% |
| Fragmented genes | 9.2% | 4.8% | 4.8% | 1.1% |
| Missing genes | 18.3% | 3.5% | 2.9% | 2.2% |
| Reference | McGaugh et al. 2015 | This study | This study | Eckalbar et al, 2013 |

**Table 4.** Annotation of *Sceloporus undulatus de novo* transcriptome assembly using 4 tissues. Unique annotation numbers between parentheses.

| **Annotation** |  |
| --- | --- |
| Annotated genes | 467,658 |
| Annotated transcript isoforms | 547,370 |
| Annotated isoforms/gene | 1.17 |
| Transcripts with Swiss-Prot annotation | (71,944) |
| Transcripts with PFAM annotation | 51,018 (46,432) |
| Transcripts with KEGG annotation | 65,694 (21,520) |
| Transcripts with GO annotation | 73,936 (66,554) |

### Genome Assembly Annotation

Using the top 24 largest scaffolds of the SceUnd1.0 assembly (we refer to this set as SceUnd1.0_top24),we used the Funannotate v1.5.0 pipeline (https://github.com/nextgenusfs/funannotate) for gene prediction and functional annotation. Funannotate uses RNAseq data and the Tetrapoda BUSCO [44] dataset to train the *ab initio* gene prediction programs Augustus [59] and GeneMark-ET [60]. Evidence Modeler is used to generate the consensus from Augustus and GeneMark-ES/ET. In the training step, we used four raw RNAseq datasets described in Table 2 that contained a total of 68 sequenced libraries. tRNAscan-SE [61] was used to predict tRNA genes. Finally the genes were functionally annotated via InterProScan [62], Eggnog [63], PFAM [54], UniProtKB [64], MEROPS [65], CAZyme, and GO ontology. We also used DIAMOND blastp [66] to compare the predicted proteins to ENSEMBL human, chicken, mouse, and gene anole lizard databases (Supplemental files: SceUnd1.0_top24.gff3; SceUnd1.0_top24_CompliedAnnotation.csv). Our annotation pipeline predicted 54,149 genes, 15,472 of which were attributed meaningful functional annotation beyond “hypothetical protein”. Through BLAST of the predicted protein coding genes we found 21,050 (39%) had hits in ENSEMBL. We then quantified the number of BUSCO genes identified in the predicted proteins from the Funannotate pipeline and found 79.1%, which corresponds to an 11.6% decrease from the number of complete BUSCO genes in the SceUnd1.0 genome assembly, which suggests this first version of annotation can be improved.

We used annotation and sequence homology to identify the X chromosome. Sex chromosomes are highly variable among *Sceloporus* species, and the genus appears to have evolved multiple XY systems independently [31]. However, some species, including *S. undulatus*, do not appear to have morphologically distinct sex chromosomes [67]. While the ancestral condition is heteromorphic chromosomes with a minute Y, many species within the genus demonstrate multiple sex chromosome heteromorphisms (i.e. multiple forms of the X chromosome) or have evolved indistinct sex chromosomes, such as the *undulatus* species group [18]. To identify the scaffold likely representing the X chromosome within *S. undulatus*, we blasted 16 X-linked genes from the green anole downloaded from Ensembl (AnoCar2.0: ACAD10, ADORA2A, ATP2A2, CCDC92, CIT, CLIP1, CUX2, DGCR8, FICD, MLEC, MLXIP, ORAI1, PLBD2, PUS1, TMEM119, ZCCHC8) [68, 69] to the SceUnd1.0. They almost exclusively map to the tenth largest scaffold, the fourth predicted microchromosome (Figures 2b, 3), indicating that it is likely the X chromosome. The Y chromosome could not be independently identified from the assembly, most likely due to the homomorphic nature of *S. undulatus* sex chromosomes; higher sequence homology may have caused the Y chromosome to assemble with the X chromosome [31].

**Figure 3.**
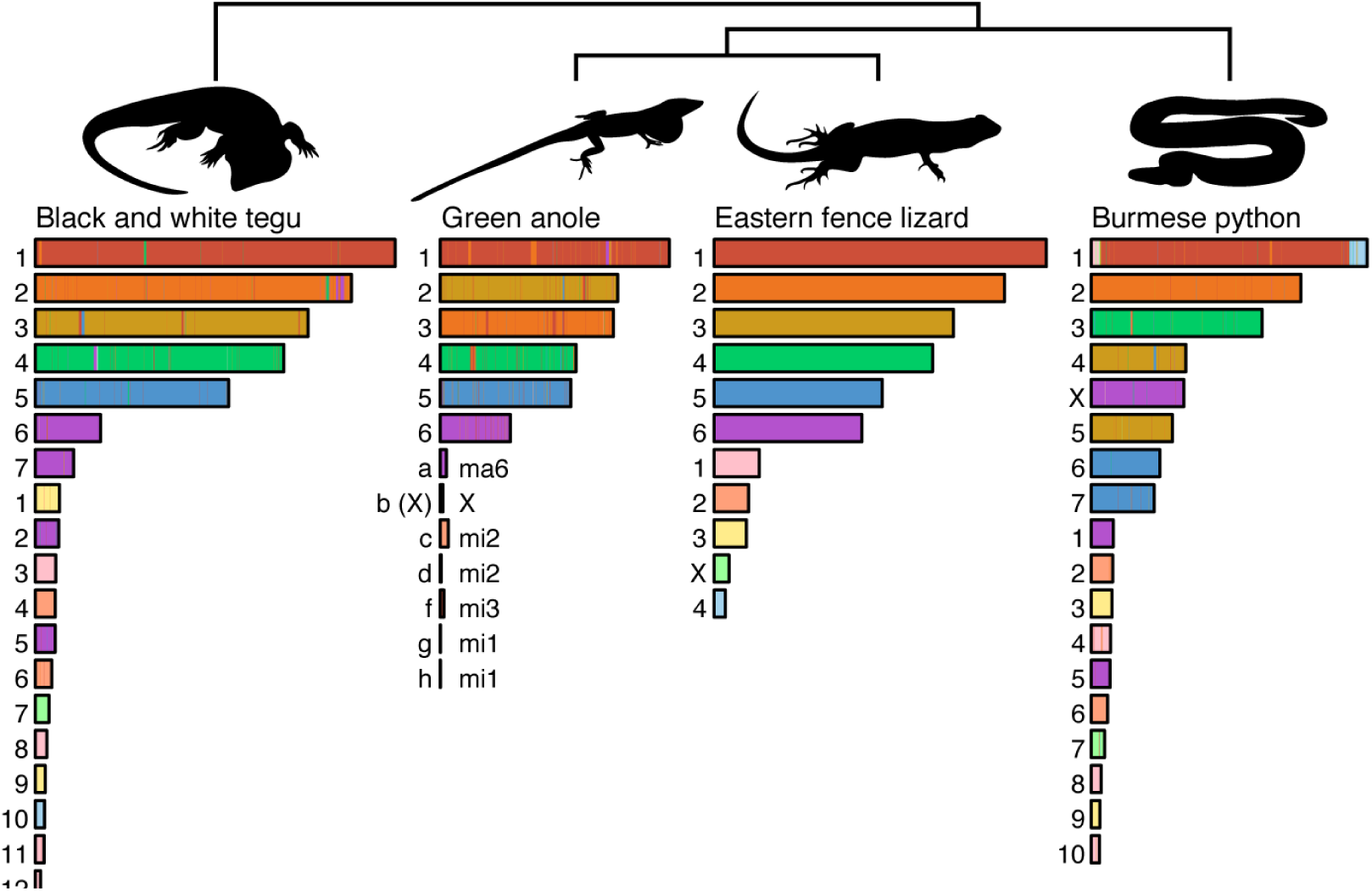
Marker-based synteny painting of fence lizard scaffolds/chromosomes onto the tegu, green anole, and python assemblies, depicted from left-to-right as tegu, green anole, fence lizard, and python. The color indicates synteny for that scaffold. The linkage groups representing macrochromosomes and microchromosomes are numbered independently for each species. Green anole linkage groups are labeled with lowercase letters, and the syntenic fence lizard chromosomes are listed to the right. Sex chromosomes are indicated with uppercase letters, where known.

### Mitochondrial Genome Assembly

The mitochondrial genome was not captured by the genome sequencing approaches, likely due to how these types of libraries are prepared. Mitochondrial sequence data obtained via RNAseq can be effectively assembled into whole mtDNA genomes [70-73]. We used RNAseq reads from 18 *S. undulatus* individuals from the RNAseq Dataset 4 (Table 2), which are from the same population as the individuals used for the genome sequencing. We used Trimmomatic v0.37 [50] to clean the raw reads and then mapped the clean reads to a complete *S. occidentalis* mtDNA genome [74] using BWA v0.7.15 [75]. Of the 632,987,330 total cleaned reads, 9.73% mapped to the *S. occidentalis* mtDNA genome with an average read depth of 5,164.42 reads per site per individual. After sorting and indexing mapped reads with SAMTOOLS v1.6 [76], we used the mpileup function in SAMTOOLS to build a consensus mitochondrial genome (mtGenome) excluding the reference and filling the no-coverage regions with “N” to generate 100% coverage of the mtGenome based on the consensus across the 18 individuals. We mapped the consensus genome to the well-annotated *Anolis carolinensis* mtGenome with MAFFT v1.3.7 [77] and transferred the annotation using the “copy annotation” command in GENEIOUS v.11.1.5 [78]. Annotations from the *A. carolinensis* mtGenome (17,223 bp) transferred well to the newly assembled *S. undulatus* mtGenome (17,072 bp), with 13 protein coding genes, 22 tRNA regions, 2 rRNA regions, and a control region (see full list in Supplemental File). The mitochondrial genome and the annotation are provided as supplemental data.

### Addressing reference assembly quality using population-level transcriptomic and genomic data

In developing the high-quality reference genome for *S. undulatus*, we produced three assemblies using increasing amounts of data, for correspondingly greater costs. To assess the utility of each of the assemblies for addressing ecological genomic questions, we use two datasets: RNAseq and whole genome resequencing.

First, we used RNAseq Dataset 4 (Table 5) from n= 18 males that were sampled from the same population (Alabama) as the individuals that were used to develop the reference assemblies; we then used these data to test whether the percentage of reads that mapped to the reference varied depending on which assembly we used as a reference. RNAseq data were cleaned with Trimmomatic v0.37 [50] and mapped with HISAT2 v2.1.0 [79] to each of the three *S. undulatus* genome assemblies. The percentage of reads that mapped were calculated using SAMTOOLS v1.6 flagstat [76]. We found negligible differences in mapping the RNAseq data to the SuperNova, HiRise and PBJelly assemblies where 81.49%, 82.37%, and 82.28% of cleaned reads mapped, respectively (Table 6).

**Table 5.** RNAseq datasets used for training in the genome annotation pipeline. Datasets 1 and 2 were used in the *de novo* transcriptome assembly.

| Data Set | Tissue | Age | Sex | Treatment/<br>Condition | Data<br>Type | NCBI SRA<br>Accession # |
| --- | --- | --- | --- | --- | --- | --- |
| **1. This Paper** | Skeletal muscle | Adult | Female | Post-reproductive | 100 bp PE | SAMN06312743 |
|  | Brain | Adult | Female | Post-reproductive | 100 bp PE | SAMN06312741 |
|  | Whole Embryo | Embryo | N/A |  | 100 bp PE | SAMN06312742 |
| **2. McGaugh**<br>**et al. 2015** | Liver | Juvenile |  | Control Lab | 100 bp PE | SRR629640 |
| **3. Cox et al.**<br>**In Review** | Liver | Juvenile | Female | Blank | 125 bp PE | SAMN14774299 |
|  | Liver | Juvenile | Male | Castrated | 125 bp PE | — |
|  | Liver | Juvenile | Male | Control | 125 bp PE | SAMN14774321 |
|  | Liver | Juvenile | Female | Testosterone | 125 bp PE |  |
|  | Liver | Juvenile | Male | Testosterone | 125 bp PE |  |
| **4. Simpson**<br>**et al. In Prep.** | Liver | Adult | Male | Control Lab | 150 bp PE | SAMN08687228 |
|  | Liver | Adult | Male | Acute Heat Stress | 150 bp PE | — |
|  | Liver | Adult | Male | Fire Ant Bitten | 150 bp PE | SAMN08687245 |

McGaugh SE, Bronikowski AM, Kuo C-H, Reding DM, Addis EA, Flagel LE, et al. Data from: Rapid molecular evolution across amniotes of the IIS/TOR network. Dryad Digital Repository. http://dx.doi.org/10.5061/dryad.vn872. 2015.
Cox, C. L., A. K. Chung, D. C. Card, T. A. Castoe, N. Pollock, H. John-Alder, and R. M. Cox. Evolutionary regulation of sex-biased gene expression and sexual dimorphism.
Simpson, D., R. Telemeco, T. Langkilde, T. S. Schwartz. Different ecological stressors have contrasting transcriptomic responses.

**Table 6.** Comparison of type of genome assembly as a reference for population-level analyses for RNAseq and Whole Genome Sequencing of individual from Alabama (AL, either low or high coverage), Tennessee (TN) and Arkansas (AR). Datasets were mapped to either the Supernova Assembly containing only the 10X Genomics data, the HiRise Assembly, or the PBJelly assembly (SceUnd1.0). Average SAMTOOLS QC-passed reads, reads mapped, and percentage of mapped QC-passed reads for every sequencing depth and population. Average whole-genome coverage and theoretical HET SNP sensitivity for every sequencing depth and population.

|  |  | **RNAseq-AL** | **Low Cov-AL** | **High Cov-AL** | **High Cov-TN** | **High Cov-AR** |
| --- | --- | --- | --- | --- | --- | --- |
| **PBJelly** | **QC-passed Reads** | 3.29E7 ± 6.84E6 | 5.09E7 ± 3.35E7 | 3.31E8 ± 2.64E7 | 3.45E8 ± 9.29E7 | 3.31E8 ± 6.09E7 |
|  | **Reads Mapped** | 2.71E7 ± 6.25E6 | 5.06E7 ± 3.33E7 | 3.29E8 ± 2.63E7 | 3.41E8 ± 9.05E7 | 3.22E8 ± 6.66E7 |
|  | **% Reads Mapped** | 82.28 ± 0.09 | 99.46 ± 0.11 | 99.47 ± 0.08 | 98.97 ± 0.61 | 97.00 ± 4.78 |
|  | **Whole-genome (X)** | NA | 3.36 ± 2.97 | 21.75 ± 11.46 | 22.04 ± 12.14 | 21.04 ± 11.64 |
|  | **HET SNP sensitivity** | NA | 0.55 | 0.88 | 0.87 | 0.86 |
| **HiRise** | **QC-passed Reads** | 3.30E7 ± 6.86E6 | 5.11E7 ± 3.36E7 | 3.33E8 ± 2.66E7 | 3.47E8 ± 9.39E7 | 3.33E8 ± 6.14E7 |
|  | **Reads Mapped** | 2.71E7 ± 6.30E6 | 5.07E7 ± 3.34E7 | 3.30E8 ± 2.65E7 | 3.43E8 ± 9.13E7 | 3.23E8 ± 6.69E7 |
|  | **% Reads Mapped** | 82.37 ± 0.09 | 99.29 ± 0.11 | 99.29 ± 0.08 | 98.80 ± 0.60 | 96.84 ± 4.75 |
|  | **Whole genome (X)** | NA | 3.56 ± 2.95 | 23.02 ± 10.52 | 23.33 ± 11.25 | 22.27 ± 10.81 |
|  | **HET SNP sensitivity** | NA | 0.58 | 0.93 | 0.91 | 0.91 |
| **SuperNova** | **QC-passed Reads** | 3.28E7 ± 6.83E6 | 5.11E7 ± 3.36E7 | 3.33E8 ± 2.66E7 | 3.47E8 ± 9.39E7 | 3.33E8 ± 6.14E7 |
|  | **Reads Mapped** | 2.68E7 ± 6.19E6 | 5.07E7 ± 3.34E7 | 3.30E8 ± 2.65E7 | 3.43E8 ± 9.13E7 | 3.23E8 ± 6.69E7 |
|  | **% Reads Mapped** | 81.49 ± 0.09 | 99.29 ± 0.11 | 99.29 ± 0.08 | 98.80 ± 0.60 | 96.84 ± 4.75 |
|  | **Whole-genome (X)** | NA | 3.56 ± 2.95 | 23.02 ± 10.52 | 23.33 ± 11.25 | 22.27 ± 10.81 |
|  | **HET SNP sensitivity** | NA | 0.58 | 0.93 | 0.91 | 0.91 |

Second, we prepared genomic DNA libraries for massively parallel sequencing for n=10 *S. undulatus* individuals (6 females, 4 males) from the same Alabama population as the individuals that were used to develop the reference assemblies. We also prepared libraries for n=5 *S. undulatus* individuals (1 female, 4 males) from Edgar Evins, Tennessee, and for n=5 individuals (2 females, 3 males) from St. Francis, Arkansas. This Arkansas population is at the boarders of the *S. undulatus* and *S. consobrinus* geographic distributions making its taxonomic status uncertain [18]. Specifically, we followed standard protocols for tissue DNA extraction from toe and/or tail clips with OMEGA EZNA Tissue spin-column kits. We then prepared sequencing libraries using the Illumina TruSeq Nano kit. We multiplexed these libraries with other individuals not included in this analysis and sequenced the library pool across two Illumina NovaSeq 6000 S4 sequencing runs. Five individuals from each of the three populations were sequenced to ∼20x average read coverage; the remaining five individuals from Alabama were sequenced to lower coverage (∼3x). Raw sequence read data were trimmed with Trimmomatic [50] and mapped separately to each of the three *S. undulatus* assemblies with bwa_mem [75] to each of the assemblies. SAMTOOLS flagstat [76] was used to calculate the total number of alignments in the .sam files generated during mapping and the number of shotgun reads that mapped to each assembly. The CollectWgsMetrics tool from the Picard Toolkit [80] was used to calculate genome-wide coverage of the mapped reads for each individual and assembly. For all sequencing depths and populations, we observed that fewer total alignments to the PBJelly Assembly than to either the HiRise or Supernova Assemblies (Table 6). Even though there were <0.5% fewer total reads that passed QC with the PBJelly Assembly/ SceUnd1.0, a higher percentage of the QC-passed reads mapped to this assembly than to either the HiRise or Supernova Assemblies (Table 6). We also determined that individuals from the same population as the *S. undulatus* individuals used to create these reference assemblies had a higher percentage of reads map to the assemblies than individuals from the Tennessee or Arkansas populations (Table 6). Those reads had lower whole-genome coverage and lower theoretical HET SNP sensitivity (i.e., sites that have increased rates of heterozygosity and might be SNPs) when mapped to the PBJelly/ SceUnd1.0 Assembly than either the HiRise or Supernova Assemblies (Table 6).

Both the RNAseq and the whole genome resequencing datasets support the conclusion that the 10X Chromium data that was used for the SuperNova Assembly covered the genome and that the HiC data (included in the HiRise Assembly) and the PacBio data (included in the final PBJelly Assembly) did not increase the amount of sequence information. Rather, the use of the HiC data and PacBio data resulted in larger scaffolds and thereby slightly increased SNP sensitivity.

### Assembly and refinement of genomic data for 34 additional *Sceloporus* species

Draft reduced representation genomes are available for 34 species within *Sceloporus* [81, 82] (phylogeny in Figure 4a). We downloaded the raw genomic reads for these 34 *Sceloporus* species from the Sequence Read Archive (Study Accession SRP041983; Table 7). Genomic resources for 33 of the species were obtained using reduced representation libraries (yielding approximately 5 Gb per species), while one species, *S. occidentalis*, was sequenced using whole genome shotgun sequencing (40.88 Gb; Table 7)[81]. To improve the draft assemblies for these 34 species, we mapped these raw reads to the final assembly, SceUnd1.0, using BWA-MEM [83]. Only the 11 longest, putative chromosome scaffolds from the SceUnd1.0 were used. The GATK version 3 [84-86] RealignerTargetCreator and IndelRealigner tools were used for local realignment, and HaplotypeCaller was used to identify insertion/deletion (INDEL) and single nucleotide polymorphism (SNP) variants. These sequence variants were separated and filtered with the SelectVariants and VariantFiltration tools using the GATK base settings. BEDTools [87] ‘genomecov’ tool was used to calculate coverage and identify regions with no coverage. We generated consensus sequences for each species by writing variants back over the reference fasta and replacing nucleotides with no coverage with “N”, using BCFtools [76] ‘consensus’ for SNPs and BEDTools ‘maskfasta’ for indels and regions with no mapping coverage (Supplemental Code File).

**Table 7.**
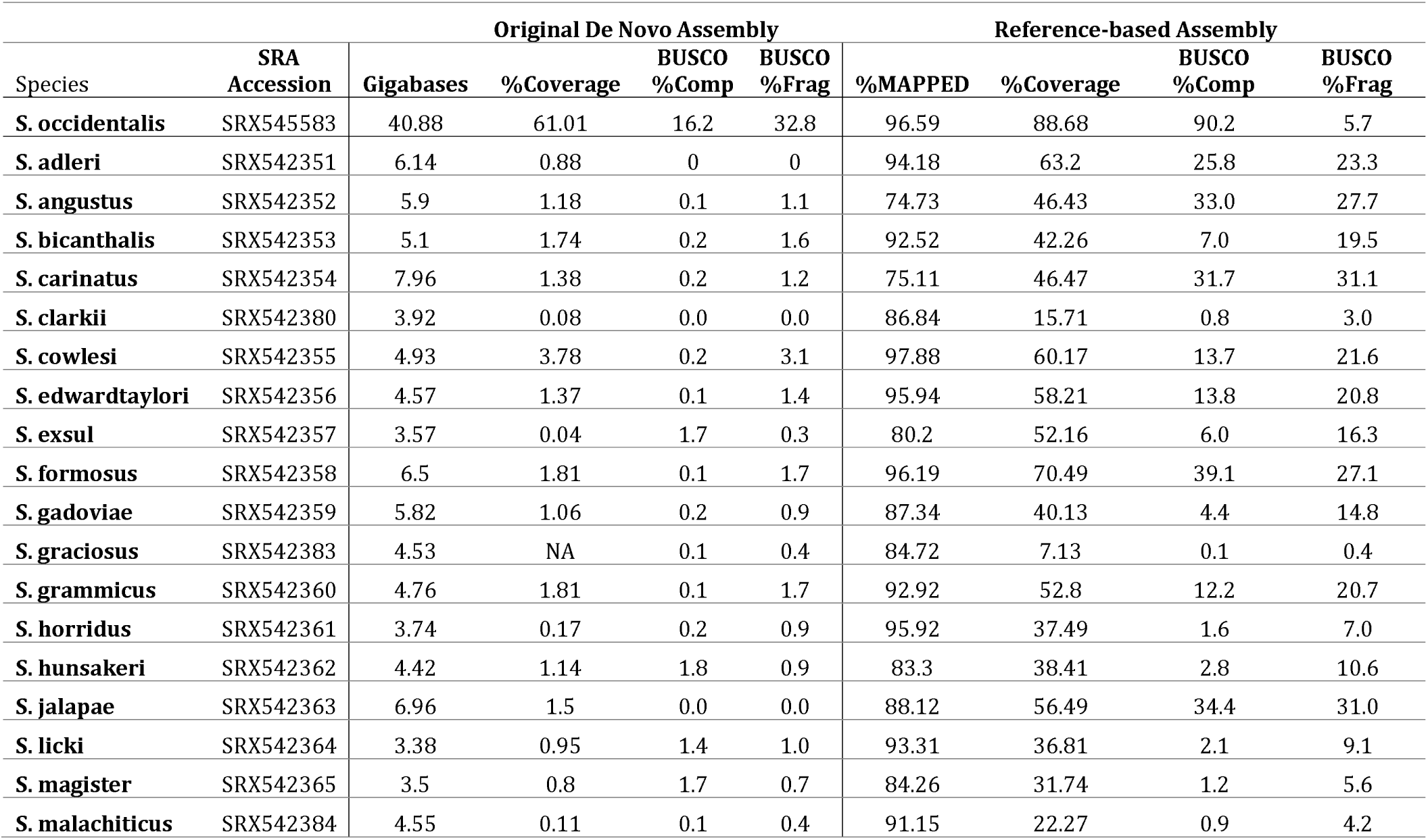
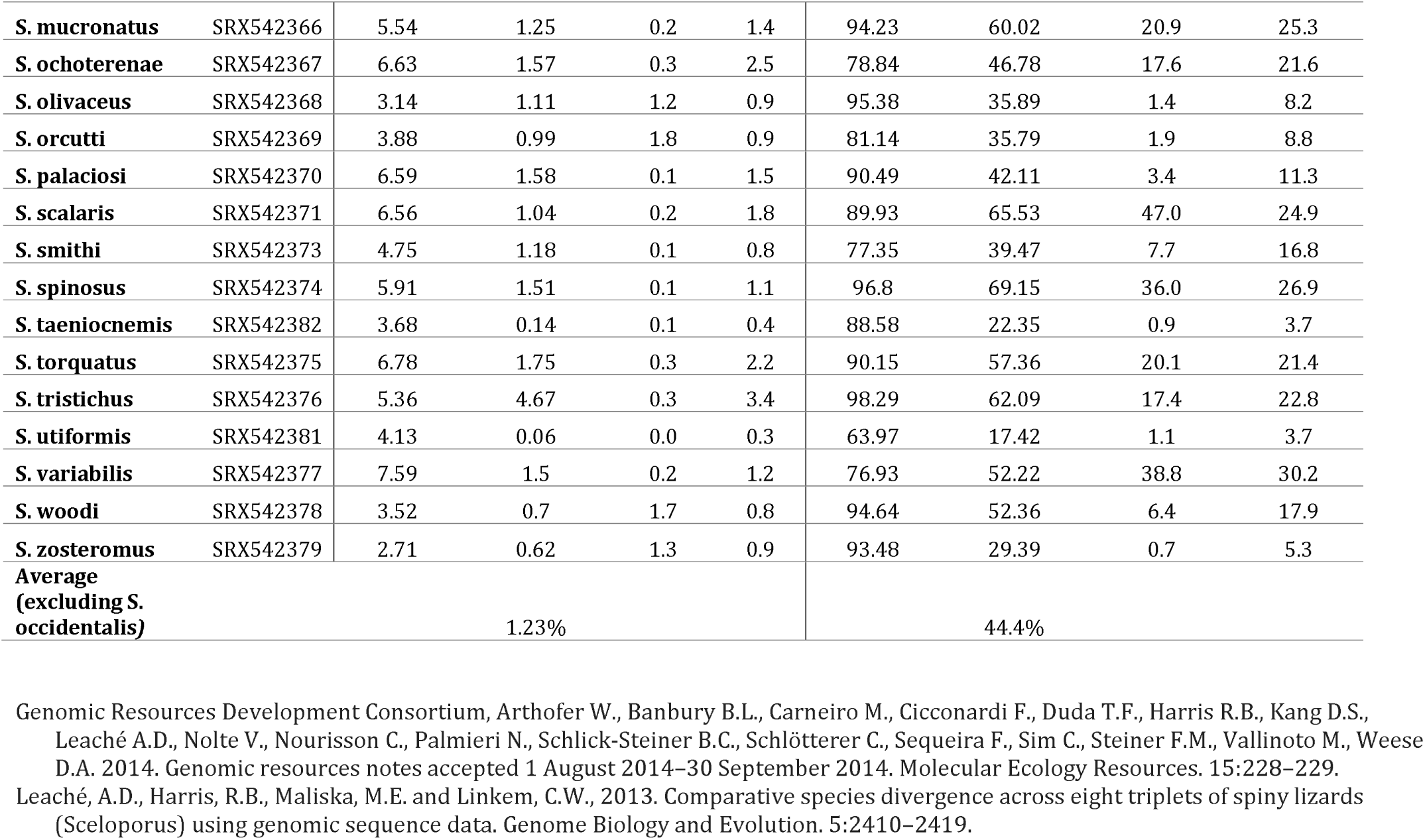
*Sceloporus* species with partial genomic sequence assemblies. Genomic resources for 34 of the species were obtained using reduced representation libraries (Arthofer et al. 2014), while one species, *S. occidentalis*, was sequenced using whole genome shotgun sequencing (Leaché et al. 2013). The data were downloaded from the Sequence Read Archive (Study Accession SRP041983; Genomic Resources Development Consortium et al., 2015).

**Figure 4.**
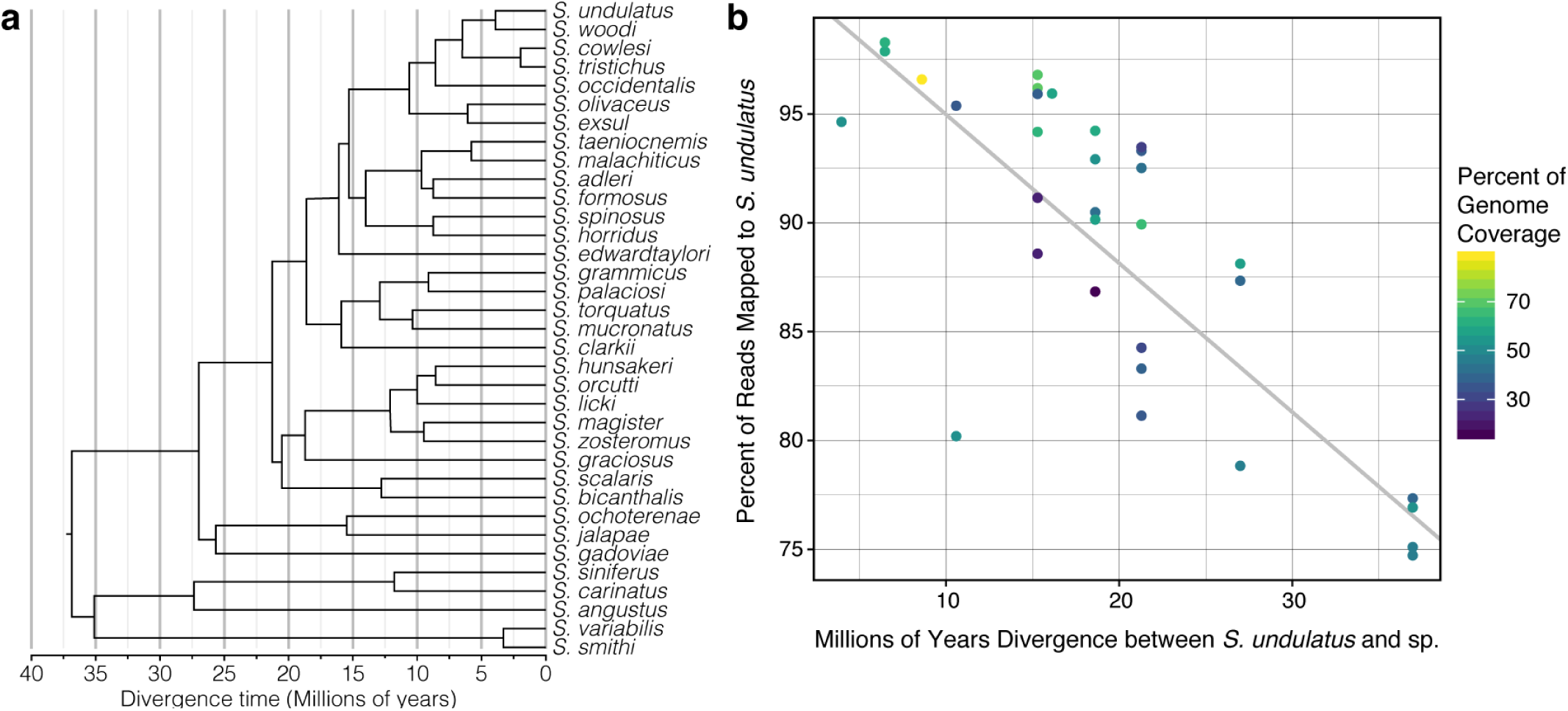
Relationship between divergence time and effectiveness of using the *Sceloporus undulatus* assembly for reference-based mapping. (a) A phylogenetic tree of *Sceloporus* species with draft genomic data. Species groups’ names are included for the groups closest to *S. undulatus*. (b) Mapping each species by % reads mapped and time of divergence from *S. undulatus* with a linear regression. The color of the dots represents the percent of the genome that is covered, which was affected by the number of redundant sequences in the reduced representation library for a particular species.

Mapping the reduced representation genome data from the 33 additional *Sceloporus* species improved the assemblies for the species. For the species with ∼5Gb of sequencing data, this improvement was from an average of 1.23% to an average of 44.4% coverage, and *S. occidentalis* with 41Gb of data improved from 61.0% to 88.7% coverage (Table 7). Across the 33 species with 5Gb of data, the BUSCO genes identified (complete and fragmented) in the reference-based assemblies ranged from 0.5 to 71.9% (complete and fragmented), whereas *S. occidentalis* had 95.9% BUSCO genes (complete and fragmented) identified, similar to our *S. undulatus* SuperNova Assembly (Table 7). Notably, across the *Sceloporus* genus, the percent of the raw data that mapped to the reference was significantly negatively correlated with divergence time to the reference *S. undulatus* (p<0.0001, r=0.779; Figure 4b). For species that are less then ∼20 million years diverged from *S. undulatus* >90% of reads mapped; the percentage of reads mapped declined to 75% when divergence was greater than 35 million years (Figure 4b).

It is important to note that the reference-based assemblies produced for these 34 species will correspond 1:1 with the synteny of the *S. undulatus* scaffolds. However, *Sceloporus* is unique among squamates for remarkable chromosome rearrangements with karyotypes ranging from 2N=22 to 2N=46 [31]. Therefore, the genome assemblies for species with karyotypes other than 2N=22 (the *S. undulatus* reference) or with large chromosomal inversions will not be reliable for addressing questions related to genomic architecture or structural variation [88]. These genome assemblies will, however, prove useful for analyses of protein and gene sequence evolution and for mapping and pseudomapping-based RNAseq analyses of gene expression across the genus to understand behavioral ecology, physiology, developmental biology, and more.

### Analysis of synteny with other squamate chromosome-level genomes

As another benchmark of genome completeness, and to generate an initial look at chromosome evolution among squamates, we performed synteny analysis of the Eastern fence lizard (*S. undulatus*) SceUnd1.0 assembly with the green anole (*Anolis carolinensis*, AnoCar2.0) and with recently published chromosome-level assemblies for the Burmese python (*Python bivittatus*) [89] and the Argentine black and white tegu lizard (*Salvator merianae*) [42] (available at https://www.dnazoo.org/). The SceUnd1.0 scaffolds representing the 11 putative chromosomes were used to produce 1000 bp-long markers excluding gapped regions. Using BLAST, these markers were compared to the predicted chromosomes from the python and tegu HiC assemblies. BLAST hits for each were filtered to only include hits that were 80% identity, at least 500bp long, and part of 4 consecutive hits from the same Eastern fence lizard chromosome. Using these results, the Eastern fence lizard chromosomes were painted onto the anole, python, and tegu chromosomes to visualize large-scale synteny (Figure 3).

From this marker-based synteny painting, we found that Eastern fence lizard has fewer chromosomes than each of the other three species, corresponding to known karyotypes for these species. Notably, many of the differences in the Eastern fence lizard relative to the other species are the result of fusion of microchromosomes (e.g. compare tegu microchromosomes 1 and 9 to Eastern fence lizard microchromosome 3) or occasionally of a microchromosome to macrochromosomes (e.g. compare tegu macrochromosomes 6 and 7 and microchromosomes 2 and 5 to the Eastern fence lizard macrochromosome 6), although the synteny of the macrochromosomes was largely conserved.

The putative sex chromosome in the SceUnd1.0 assembly (Figure 3) is syntenic to the anole X chromosome, and a microchromosome in each of the other two squamates. However, it is not syntenic to the python X chromosome, which is syntenic to the Z chromosome in other snakes. The tegu sex chromosome has not been identified.

## Discussion

For the advancement of reptilian genomic and transcriptomic resources, we provide a high-quality, chromosome-level genome assembly for the Eastern fence lizard, *Sceloporus undulatus, de novo* transcriptomes for *S. undulatus* encompassing multiple tissues and life stages, and improved draft genome assemblies from 34 additional *Sceloporus* species. In the final reference assembly, SceUnd1.0, the largest 11 scaffolds contain 92.6% (1.765 of 1.905 Gb) of the genome sequence; these 11 scaffolds likely represent the 6 macro- and 5 microchromosomes of *S. undulatus*, based on karyotype, genome size, BUSCO analysis, and synteny with other squamate genomes. The remaining small scaffolds may contain some chromosome segments that could not be assembled, misassembled regions, and/or duplicated genes.

In comparing the three levels of reference genome assemblies, we found that the first level using only the 10X Genomics and the SuperNova Assembly contained all, or very nearly all, of the protein-coding regions of the genome within its contigs (based on BUSCO and mapping of RNAseq and whole genome resequencing data). By including the Hi-C data, the contiguity of the HiRise Assembly dramatically improved, joining contigs into chromosome-length scaffolds, but had minimal effect on mapping percentages for either RNAseq or WGS. The inclusion of the PacBio data in the final PBJelly Assembly to produce SceUnd1.0 closed some gaps but yielded a relatively small improvement after the already dramatic improvements from the Hi-C data.

While it is now becoming possible to obtain a reference genome assembly for almost any organism, the quality and cost of reference genome assemblies vary considerably depending on the technologies used. This presents researchers with an important question: what levels of sequencing effort and assembly quality are required for a particular ecological genomics study? Important factors that must be considered include the sequencing depth, sequence contiguity, and thoroughness of annotation. Our study demonstrates that the SuperNova Assembly was sufficient for mapping RNAseq and whole genome resequencing, while the more expensive assemblies (HiRise and PBJelly) were necessary to achieve high-level continuity and chromosome-level scaffolding.

Genome assemblies of high-quality and contiguity are critical for understanding organismal biology in a wide range of contexts that includes behavior, physiology, ecology, and evolution, on scales ranging from populations to higher-level clades. From RNAseq to ChIP-seq and epigenetics, large-scale sequencing is rapidly becoming commonplace in ecological genomics to address fundamental questions of how organisms directly respond to their environment and how populations evolve in response to environmental variation. Many advanced molecular tools are typically reserved for traditional model organisms but with the large foundation of ecological and physiological data available for *S. undulatus*, a high-quality reference genome opens the door for these molecular techniques to be used in this ecological model organism. For example, with the recent demonstration of CRISPR-Cas9 gene modification in a lizard, the brown anole [90], a genome reference will facilitate the application of gene drive technologies for functional genomic studies in *Sceloporus* lizards. This reference will provide a foundation for whole genome studies to understand speciation and hybridization among closely related species utilizing low coverage re-sequencing, or as a point of comparison with more distantly related species relative to the chromosomal inversions and large-scale genome architectural changes common in the clade. *Sceloporus undulatus* and other lizards in the genus *Sceloporus* exhibit evolutionary reversals in sexual size dimorphism and dichromatism and they have been used to demonstrate that androgens such as testosterone can inhibit growth in species (such as *S. undulatus*) in which females are the larger sex [19, 91-93]. This SceUnd1.0 chromosome-level genome assembly would support ChIPseq or *in silico* analyses to identify sex hormone response elements. In addition, this assembly will facilitate the identification of signatures of exposure to environmental stressors in both gene expression and epigenetic modification [94] to evaluate pressing questions on how climate change and invasive species affect local fauna. All of these uses for a chromosome-level genome assembly provide valuable extensions to ongoing work in the *Sceloporus* genus.

## Availability of Supporting Data

1. All three genome assemblies are provided as supplemental data
  a. SuperNova assembly containing data from 10X Genomics Chromium: GenomeAssembly_SuperNova_Sceloporus_undulatus_pseudohap.fasta.gz
  b. HiRise assembly containing the 10X Genomics data with the addition of the Hi-C data: GenomeAssembly_HiRise_Sceloporus_undulatus.fasta.gz
  c. PBJelly Assembly (SceUnd1.0) containing the 10X Genomics data, the Hi-C data, with the addition of PacBio data: GenomeAssembly_SceUnd1.0_PBJELLY.fasta.gz
2. Tissue-Embryo Transcriptomes and annotation are provided as supplemental data.
  a. Transcriptome File: TranscriptomeAssembly_Tissues-Embryo_Trinity.fasta
  b. Annotation File: TranscriptomeAssembly_Tissues-Embryo_Transdecoder.gff3
3. Truncated assembly used for annotation pipeline (SceUnd1.0_top24)
  a. SceUnd1.0_top24.fasta. This file contains only the longest 24 scaffolds and they have been renamed 1-24 from longest to shortest.
  b. Funannotate Folder: contains that annotation files
  c. SceUnd1.0_top24_CompliedAnnotation.csv
4. The mitochondrial genomes and the annotation are provided as supplemental data.
  a. MitoGenomeAssembly_Sceloporus_undulatus.fasta
  b. MitoGenomeAssembly_Sceloporus_undulatus_Annotation.gff
5. The reference-based assemblies for the 34 *Sceloporus* species.
  a. GenomeAssemblies_34Sceloporus.tar.gz
  b. Code for generated consensus sequences for each species: mkgenome_AW-AC.sh

## Acknowledgements

We are grateful for the support of the DoveTail Genomics and Auburn University Office of Information Technology and Hopper High-Performance Computing Cluster for assistance with this work. We thank Kirsty MacLeod for catching the adult male used for sequencing, sequencing and Juan Rodriguez for bioinformatic assistance.

## Authors’ Contributions

**AW:** Data curation; Formal analysis; Investigation; Validation; Visualization; Writing – original; Writing – review & editing

**RST:** Conceptualization; Data curation; Formal analysis; Investigation; Validation; Visualization; Writing – review & editing

**MBG**: Data curation; Formal analysis; Investigation; Validation; Visualization; Writing – original; Writing – review & editing

**DSW:** Data curation; Formal analysis; Software; Validation; Visualization; Writing – original; Writing – review & editing

**DYS:** Formal analysis; Software; Writing – original; Writing – review & editing

**RK:** Methodology, Formal analysis, Writing- original draft, Writing- review & editing

**AC:** Data curation; Formal analysis; Methodology; Software, Validation, Visualization

**APS:** Formal analysis; Writing – original draft; Writing – review & editing

**CLC:** Conceptualization; Data Curation; Investigation; Funding Acquisition; Writing-review & editing

**GP:** Funding acquisition; Supervision, Writing – review & editing.

**MT:** Data curation; Formal analysis; Methodology; Funding acquisition; Writing – review & editing

**TL:** Conceptualization; Funding acquisition; Resources; Writing – review & editing

**KK:** Conceptualization; Funding acquisition; Resources; Writing – review & editing

**MWS:** Resources; Funding Acquisition; Writing-review & editing

**ADL:** Conceptualization; Data curation; Funding acquisition; Methodology; Writing – original; Writing – review & editing

**MJA:** Conceptualization; Funding acquisition; Writing – review & editing

**MEG:** Conceptualization; Writing – review & editing

**HJA:** Investigation; Funding acquisition; Writing – review & editing

**RMC:** Conceptualization; Funding acquisition; Investigation; Writing – review & editing **TSS:** Conceptualization; Data curation; Funding acquisition; Investigation; Project Administration; Resources; Supervision; Writing – original; Writing – review & editing.

All authors have read and approved the final version of the manuscript.

## Supplementary Methods and Results

### Full list of genes identified in the mitochondrial genome

Annotations from the *A. carolinensis* mitochondrial genome (17,223 bp) transferred well to the newly assembled *S. undulatus* mitochondrial genome (17,072 bp), with 13 protein coding genes (ATP6, ATP8, COX1, COX2, COX3, CYTB, ND1, ND2, ND3, ND4, ND4L, ND5, ND6), 22 tRNA regions (tRNA-Phe, tRNA-Val, tRNA-Leu, tRNA-Ile, tRNA-Gln, tRNA-Met, tRNA-Trp, tRNA-Ala, tRNA-Asn, tRNA-Cys, tRNA-Tyr, tRNA-Ser, tRNA-Asp, tRNA-Lys, tRNA-Gly, tRNA-Arg, tRNA-His, tRNA-Ser, tRNA-Leu, tRNA-Glu, tRNA-Thr, tRNA-Pro), 2 rRNA regions (12S, 16S), and a control region.

**Table S1.** Contig length statistics for *Sceloporus undulatus de novo* transcriptome assemblies. 4 tissues = 3 tissues (brain, skeletal muscle and embryos) + 1 tissue (liver; McGaugh et al, 2015).

|  | 1 tissue | 3 tissues | 4 tissues |
| --- | --- | --- | --- |
| Minimum length | 201.0 | 201.0 | 201.0 |
| 1<sup>st</sup> Quartile | 266.0 | 266.0 | 266.0 |
| Median | 382.0 | 377.0 | 375.0 |
| Mean | 829.9 | 822.4 | 781.0 |
| 3<sup>rd</sup> Quartile | 808.0 | 732.0 | 711.0 |
| Maximum length | 16,776.0 | 30,410.0 | 30,258.0 |

**Table S2.** Reads mapped to *Sceloporus undulatus de novo* transcriptome assembly using 4 tissues.

| Read classification | Counts | Percentage of mapped reads |
| --- | --- | --- |
| Proper pairing | 170,981,981 | 97.10% |
| Left read only | 3,778,790 | 2.15% |
| Right read only | 1,015,874 | 0.58% |
| Improper pairing | 310,142 | 0.18% |

**Table S3.** Representation of full-length reconstructed protein-coding genes in *Sceloporus undulatus de novo* transcriptome, using the protein set of *Anolis carolinensis* (AnoCar2.0, Ensembl) as a reference.

| Alignment coverage | Counts | Cumulative counts |
| --- | --- | --- |
| 100% | 9,874 | 9,874 |
| 90% | 1,349 | 11,223 |
| 80% | 799 | 12,022 |
| 70% | 757 | 12,779 |
| 60% | 725 | 13,504 |
| 50% | 577 | 14,081 |
| 40% | 463 | 14,544 |
| 30% | 455 | 14,999 |
| 20% | 358 | 15,357 |
| 10% | 97 | 15,454 |

**Figure S1.**
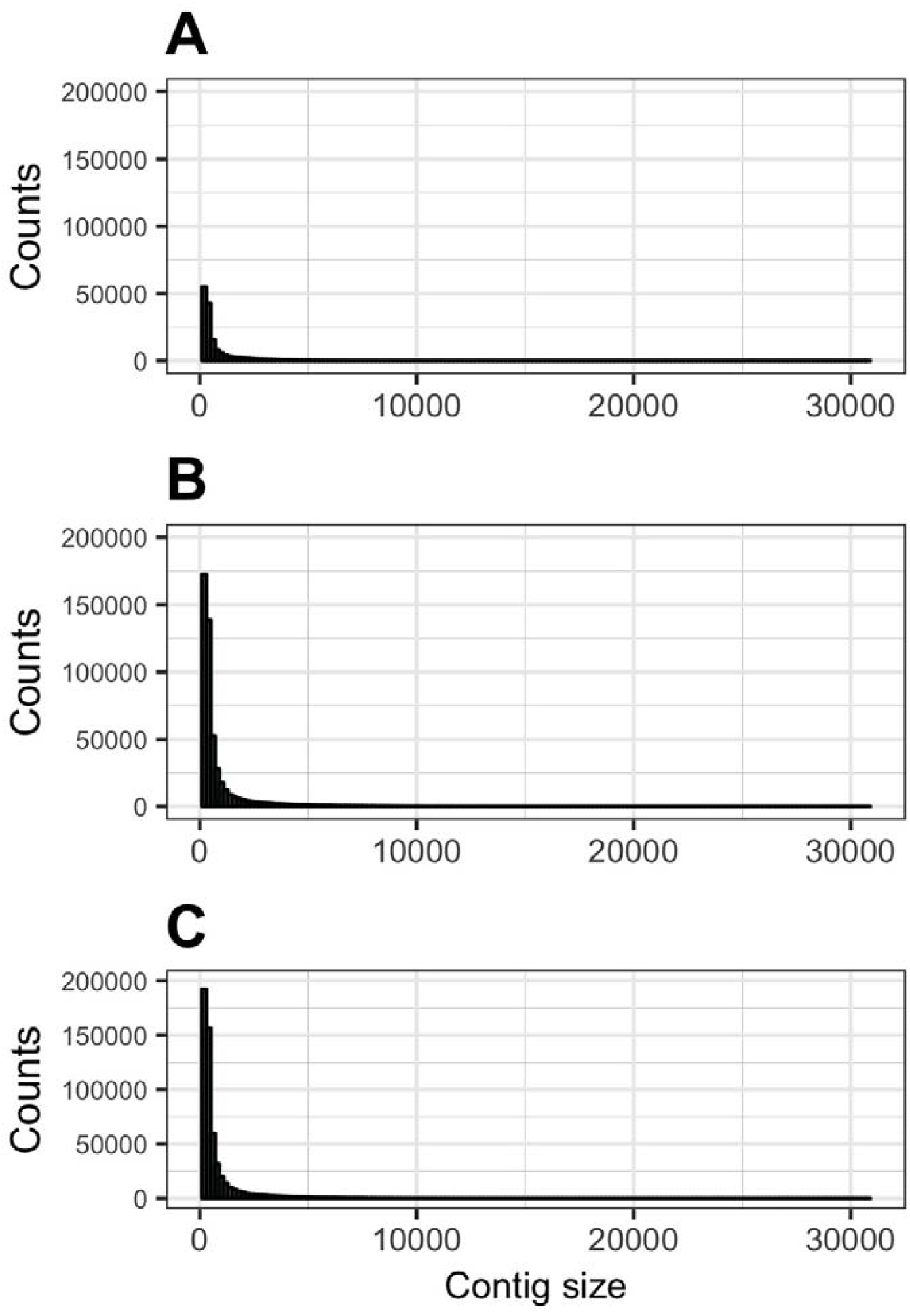
Contig sizes for different *Sceloporus undulatus* transcriptome assemblies. Assemblies used (A) the previously published single tissue transcriptome (liver [23]), (B) transcriptomes from the 3 tissues sequenced in this study (brain, skeletal muscle and embryos), and (C) the combined data set of 4 tissues ([23] and this study).

## References

1. Seebacher F. A review of thermoregulation and physiological performance in reptiles: what is the role of phenotypic flexibility? Journal of Comparative Physiology B: Biochemical, Systemic, and Environmental Physiology. 2005;175 7:453–61.

2. Kearney M, Fujita MK and Ridenour J. Lost Sex in the Reptiles: Constraints and Correlations. In: Schön I, Martens K and Dijk P, editors. Lost Sex: The Evolutionary Biology of Parthenogenesis. Dordrecht: Springer Netherlands; 2009. p. 447–74.

3. Van Dyke JU, Brandley MC and Thompson MB. The evolution of viviparity: molecular and genomic data from squamate reptiles advance understanding of live birth in amniotes. Reproduction. 2014;147 1:R15–26. doi:10.1530/REP-13-0309.

4. Rhen T and Schroeder A. Molecular mechanisms of sex determination in reptiles. Sex Dev. 2010;4 1-2:16–28. doi:10.1159/000282495.

5. Sarre SD, Ezaz T and Georges A. Transitions between sex-determining systems in reptiles and amphibians. Annu Rev Genomics Hum Genet. 2011;12:391–406. doi:10.1146/annurev-genom-082410-101518.

6. Bergmann PJ and Morinaga G. The convergent evolution of snake-like forms by divergent evolutionary pathways in squamate reptiles. Evolution. 2019;73 3:481–96. doi:10.1111/evo.13651.

7. Liu Y, Zhou Q, Wang Y, Luo L, Yang J, Yang L, et al. Gekko japonicus genome reveals evolution of adhesive toe pads and tail regeneration. Nat Commun. 2015;6:10033. doi:10.1038/ncomms10033.

8. Andrew AL, Perry BW, Card DC, Schield DR, Ruggiero RP, McGaugh SE, et al. Growth and stress response mechanisms underlying post-feeding regenerative organ growth in the Burmese python. BMC Genomics. 2017;18 1:338. doi:10.1186/s12864-017-3743-1.

9. Janes DE, Organ CL and Fujita MK. Genome evolution in Reptilia, the sister group of mammals. Annual review of genomics and human genetics. 2010;11:239–64. doi:10.1146/annurev-genom-082509-141646.

10. Alfoldi J, Di Palma F, Grabherr M, Williams C, Kong L, Mauceli E, et al. The genome of the green anole lizard and a comparative analysis with birds and mammals. Nature. 2011;477 7366:587–91. doi:10.1038/nature10390.

11. Georges A, Li Q, Lian J, O’Meally D, Deakin J, Wang Z, et al. High-coverage sequencing and annotated assembly of the genome of the Australian dragon lizard Pogona vitticeps. Gigascience. 2015;4:45. doi:10.1186/s13742-015-0085-2.

12. Xiong Z, Li F, Li Q, Zhou L, Gamble T, Zheng J, et al. Draft genome of the leopard gecko, Eublepharis macularius. Gigascience. 2016;5 1:47. doi:10.1186/s13742-016-0151-4.

13. Lind AL, Lai YYY, Mostovoy Y, Holloway AK, Iannucci A, Mak ACY, et al. Genome of the Komodo dragon reveals adaptations in the cardiovascular and chemosensory systems of monitor lizards. Nat Ecol Evol. 2019;3 8:1241–52. doi:10.1038/s41559-019-0945-8.

14. Olmo E. Trends in the evolution of reptilian chromosomes. Integr Comp Biol. 2008;48 4:486–93. doi:10.1093/icb/icn049.

15. Hedges SB, Marin J, Suleski M, Paymer M and Kumar S. Tree of life reveals clock-like speciation and diversification. Mol Biol Evol. 2015;32 4:835–45. doi:10.1093/molbev/msv037.

16. Zhang G, Li C, Li Q, Li B, Larkin DM, Lee C, et al. Comparative genomics reveals insights into avian genome evolution and adaptation. Science. 2014;346 6215:1311. doi:10.1126/science.1251385.

17. Pasquesi GIM, Adams RH, Card DC, Schield DR, Corbin AB, Perry BW, et al. Squamate reptiles challenge paradigms of genomic repeat element evolution set by birds and mammals. Nat Commun. 2018;9 1:2774. doi:10.1038/s41467-018-05279-1.

18. Leaché AD. Species Tree Discordance Traces to Phylogeographic Clade Boundaries in North American Fence Lizards (Sceloporus). Systematic Biology. 2009;58 6:547–59. doi:10.1093/sysbio/syp057.

19. John-Alder HB, Cox RM, Haenel GJ and Smith LC. Hormones, performance and fitness: Natural history and endocrine experiments on a lizard (Sceloporus undulatus). Integr Comp Biol. 2009;49 4:393–407. doi:10.1093/icb/icp060.

20. Buckley LB, Urban MC, Angilletta MJ, Crozier LG, Rissler LJ and Sears MW. Can mechanism inform species’ distribution models? Ecology Letters. 2010;13 8:1041–54.

21. Warner DA and Andrews RM. Nest-Site Selection in Relation to Temperature and Moisture by the Lizard Sceloporus undulatus. Herpetologica. 2002;58 4:399–407. doi:10.1655/0018-0831(2002)058[0399:Nsirtt]2.0.Co;2.

22. Telemeco RS, Fletcher B, Levy O, Riley A, Rodriguez-Sanchez Y, Smith C, et al. Lizards fail to plastically adjust nesting behavior or thermal tolerance as needed to buffer populations from climate warming. Glob Chang Biol. 2016; doi:10.1111/gcb.13476.

23. Blackburn DG, Gavelis GS, Anderson KE, Johnson AR and Dunlap KD. Placental specializations of the mountain spiny lizard Sceloporus jarrovi. Journal of morphology. 2010;271 10:1153–75. doi:10.1002/jmor.10860.

24. Anderson KE, Blackburn DG and Dunlap KD. Scanning electron microscopy of the placental interface in the viviparous lizard Sceloporus jarrovi (Squamata: Phrynosomatidae). Journal of morphology. 2011;272 4:465–84. doi:10.1002/jmor.10925.

25. Lambert SM and Wiens JJ. Evolution of viviparity: a phylogenetic test of the cold-climate hypothesis in phrynosomatid lizards. Evolution. 2013;67 9:2614–30. doi:10.1111/evo.12130.

26. Angilletta JMichael J, Niewiarowski Peter H, Dunham Arthur E, Leaché Adam D and Porter Warren P. Bergmann’s Clines in Ectotherms: Illustrating a Life-History Perspective with Sceloporine Lizards. The American Naturalist. 2004;164 6:E168–E83. doi:10.1086/425222.

27. Angilletta MJ, Oufiero CE and Leaché AD. Direct and Indirect Effects of Environmental Temperature on the Evolution of Reproductive Strategies: An Information-Theoretic Approach. American Naturalist. 2006;168 4:E123–E35.

28. Tinkle DW and Ballinger RE. Sceloporus undulatus: A Study of the Intraspecific Comparative Demography of a Lizard. Ecology. 1972;53 4:570–84.

29. Lawing AM, Polly PD, Hews DK and Martins EP. Including Fossils in Phylogenetic Climate Reconstructions: A Deep Time Perspective on the Climatic Niche Evolution and Diversification of Spiny Lizards (Sceloporus). Am Nat. 2016;188 2:133–48. doi:10.1086/687202.

30. Rosenblum EB, Parent CE, Diepeveen ET, Noss C and Bi K. Convergent Phenotypic Evolution despite Contrasting Demographic Histories in the Fauna of White Sands. The American Naturalist. 2017;190 S1:S44–S56. doi:10.1086/692138.

31. Leaché AD and Sites JW. Chromosome evolution and diversification in north american spiny lizards (Genus Sceloporus). Cytogenetic and Genome Research. 2010;127 2-4:166–81. doi:10.1159/000293285.

32. Leache A and Reeder TW. Molecular Systematics of the Eastern Fence Lizard (Sceloporus undulatus): A Comparison of Parsimony, Likelihood, and Bayesian Approaches. Systematic Biology. 2002;51 1:44–68.

33. Cox RM, Butler MA and John-Alder HB. The evolution of sexual size dimorphism in reptiles. Sex, Size and Gender Roles. 2007. p. 38–49.

34. Pollock NB, Feigin S, Drazenovic M and John-Alder HB. Sex hormones and the development of sexual size dimorphism: 5alpha-dihydrotestosterone inhibits growth in a female-larger lizard (Sceloporus undulatus). J Exp Biol. 2017;220 Pt 21:4068–77. doi:10.1242/jeb.166553.

35. Trompeter WP and Langkilde T. Invader danger: Lizards faced with novel predators exhibit an altered behavioral response to stress. Hormones and Behavior. 2011;60 2:152–8. doi: http://dx.doi.org/10.1016/j.yhbeh.2011.04.001.

36. Graham SP, Freidenfelds NA, Thawley CJ, Robbins TR and Langkilde T. Are Invasive Species Stressful? The Glucocorticoid Profile of Native Lizards Exposed to Invasive Fire Ants Depends on the Context. Physiol Biochem Zool. 2017;90 3:328–37. doi:10.1086/689983.

37. Gifford ME, Robinson CD and Clay TA. The influence of invasive fire ants on survival, space use, and patterns of natural selection in juvenile lizards. Biological Invasions. 2017;19 5:1461–9. doi:10.1007/s10530-017-1370-z.

38. Angilletta MJ, Jr., Zelic MH, Adrian GJ, Hurliman AM and Smith CD. Heat tolerance during embryonic development has not diverged among populations of a widespread species (Sceloporus undulatus). Conserv Physiol. 2013;1 1:cot018. doi:10.1093/conphys/cot018.

39. Buckley LB, Ehrenberger JC, Angilletta MJ and Wilson R. Thermoregulatory behaviour limits local adaptation of thermal niches and confers sensitivity to climate change. Functional Ecology. 2015;29 8:1038–47. doi:10.1111/1365-2435.12406.

40. Carlo MA, Riddell EA, Levy O and Sears MW. Recurrent sublethal warming reduces embryonic survival, inhibits juvenile growth, and alters species distribution projections under climate change. Ecol Lett. 2018;21 1:104–16. doi:10.1111/ele.12877.

41. Zheng GX, Lau BT, Schnall-Levin M, Jarosz M, Bell JM, Hindson CM, et al. Haplotyping germline and cancer genomes with high-throughput linked-read sequencing. Nat Biotechnol. 2016;34 3:303–11. doi:10.1038/nbt.3432.

42. Roscito JG, Sameith K, Pippel M, Francoijs KJ, Winkler S, Dahl A, et al. The genome of the tegu lizard Salvator merianae: combining Illumina, PacBio, and optical mapping data to generate a highly contiguous assembly. Gigascience. 2018;7 12 doi:10.1093/gigascience/giy141.

43. Cole CJ. Chromosome Variation in North American Fence Lizards (Genus Sceloporus; undulatus Species Group). Systematic Biology. 1972;21 4:357–63. doi:10.1093/sysbio/21.4.357.

44. Simao FA, Waterhouse RM, Ioannidis P, Kriventseva EV and Zdobnov EM. BUSCO: assessing genome assembly and annotation completeness with single-copy orthologs. Bioinformatics. 2015;31 19:3210–2. doi:10.1093/bioinformatics/btv351.

45. Waterhouse RM, Seppey M, Simao FA, Manni M, Ioannidis P, Klioutchnikov G, et al. BUSCO applications from quality assessments to gene prediction and phylogenomics. Mol Biol Evol. 2017; doi:10.1093/molbev/msx319.

46. Fisher RE, Geiger LA, Stroik LK, Hutchins ED, George RM, Denardo DF, et al. A histological comparison of the original and regenerated tail in the green anole, Anolis carolinensis. Anat Rec (Hoboken). 2012;295 10:1609–19. doi:10.1002/ar.22537.

47. Ritzman TB, Stroik LK, Julik E, Hutchins ED, Lasku E, Denardo DF, et al. The gross anatomy of the original and regenerated tail in the green anole (Anolis carolinensis). Anat Rec (Hoboken). 2012;295 10:1596–608. doi:10.1002/ar.22524.

48. McGaugh SE, Bronikowski AM, Kuo C-H, Reding DM, Addis EA, Flagel LE, et al. Rapid molecular evolution across amniotes of the IIS/TOR network. Proceedings of the National Academy of Sciences. 2015;112 22:7055–60. doi:10.1073/pnas.1419659112.

49. McGaugh SE, Bronikowski AM, Kuo C-H, Reding DM, Addis EA, Flagel LE, et al. Data from: Rapid molecular evolution across amniotes of the IIS/TOR network. Dryad Digital Repository. http://dx.doi.org/10.5061/dryad.vn872. 2015.

50. Bolger AM, Lohse M and Usadel B. Trimmomatic: a flexible trimmer for Illumina sequence data. Bioinformatics. 2014;30 doi:10.1093/bioinformatics/btu170.

51. Grabherr MG, Haas BJ, Yassour M, Levin JZ, Thompson DA, Amit I, et al. Full-length transcriptome assembly from RNA-Seq data without a reference genome. Nat Biotech. 2011;29 7:644–52. doi:http://www.nature.com/nbt/journal/v29/n7/abs/nbt.1883.html#supplementary-information.

52. Huang X, Chen XG and Armbruster PA. Comparative performance of transcriptome assembly methods for non-model organisms. BMC Genomics. 2016;17:523. doi:10.1186/s12864-016-2923-8.

53. Wu CH, Apweiler R, Bairoch A, Natale DA, Barker WC, Boeckmann B, et al. The Universal Protein Resource (UniProt): an expanding universe of protein information. Nucleic Acids Res. 2006;34 doi:10.1093/nar/gkj161.

54. Finn RD, Coggill P, Eberhardt RY, Eddy SR, Mistry J, Mitchell AL, et al. The Pfam protein families database: towards a more sustainable future. Nucleic Acids Research. 2016;44 D1:D279–D85. doi:10.1093/nar/gkv1344.

55. Langmead B and Salzberg SL. Fast gapped-read alignment with Bowtie 2. Nat Methods. 2012;9 doi:10.1038/nmeth.1923.

56. Camacho C, Coulouris G, Avagyan V, Ma N, Papadopoulos J, Bealer K, et al. BLAST+: architecture and applications. BMC Bioinformatics. 2009;10:421. doi:10.1186/1471-2105-10-421.

57. Eddy SR. A new generation of homology search tools based on probabilistic inference. Genome Inform. 2009;23.

58. Eckalbar WL, Hutchins ED, Markov GJ, Allen AN, Corneveaux JJ, Lindblad-Toh K, et al. Genome reannotation of the lizard Anolis carolinensis based on 14 adult and embryonic deep transcriptomes. BMC Genomics. 2013;14 1:49. doi:10.1186/1471-2164-14-49.

59. Stanke M, Schoffmann O, Morgenstern B and Waack S. Gene prediction in eukaryotes with a generalized hidden Markov model that uses hints from external sources. BMC Bioinformatics. 2006;7:62. doi:10.1186/1471-2105-7-62.

60. Lomsadze A, Burns PD and Borodovsky M. Integration of mapped RNA-Seq reads into automatic training of eukaryotic gene finding algorithm. Nucleic Acids Res. 2014;42 15:e119. doi:10.1093/nar/gku557.

61. Lowe TM and Chan PP. tRNAscan-SE On-line: integrating search and context for analysis of transfer RNA genes. Nucleic Acids Res. 2016;44 W1:W54–7. doi:10.1093/nar/gkw413.

62. Jones P, Binns D, Chang HY, Fraser M, Li W, McAnulla C, et al. InterProScan 5: genome-scale protein function classification. Bioinformatics. 2014;30 9:1236–40. doi:10.1093/bioinformatics/btu031.

63. Huerta-Cepas J, Szklarczyk D, Forslund K, Cook H, Heller D, Walter MC, et al. eggNOG 4.5: a hierarchical orthology framework with improved functional annotations for eukaryotic, prokaryotic and viral sequences. Nucleic Acids Res. 2016;44 D1:D286–93. doi:10.1093/nar/gkv1248.

64. Bateman A, Martin MJ, O’Donovan C, Magrane M, Alpi E, Antunes R, et al. UniProt: the universal protein knowledgebase. Nucleic Acids Research. 2017;45 D1:D158–D69. doi:10.1093/nar/gkw1099.

65. Rawlings ND, Barrett AJ, Thomas PD, Huang X, Bateman A and Finn RD. The MEROPS database of proteolytic enzymes, their substrates and inhibitors in 2017 and a comparison with peptidases in the PANTHER database. Nucleic Acids Res. 2018;46 D1:D624–D32. doi:10.1093/nar/gkx1134.

66. Buchfink B, Xie C and Huson DH. Fast and sensitive protein alignment using DIAMOND. Nature Methods. 2015;12 1:59–60. doi:10.1038/nmeth.3176.

67. Sites JW, Archie JW, Cole CJ and Villela OF. A Review of Phylogenetic Hypotheses for Lizards of the Genus Sceloporus (Phrynosomatidae) – Implications for Ecological and Evolutionary Studies. Bulletin of the American Museum of Natural History. 1992; 213:1–110.

68. Rovatsos M, Altmanová M, Pokorná M and Kratochvíl L. Conserved sex chromosomes across adaptively radiated Anolis lizards. Evolution. 2014;68 7:2079–85. doi:10.1111/evo.12357.

69. Rovatsos M, Altmanová M, Pokorná MJ and Kratochvíl L. Novel X-Linked Genes Revealed by Quantitative Polymerase Chain Reaction in the Green Anole, Anolis carolinensis. G3. 2014;4 11:2107–13. doi:10.1534/g3.114.014084.

70. Smith DR. RNA-Seq data: a goldmine for organelle research. Brief Funct Genomics. 2013;12 5:454–6. doi:10.1093/bfgp/els066.

71. Schwartz TS, Arendsee ZW and Bronikowski AM. Mitochondrial divergence between slow- and fast-aging garter snakes. Exp Gerontol. 2015;71:135–46. doi:10.1016/j.exger.2015.09.004.

72. Tian Y and Smith DR. Recovering complete mitochondrial genome sequences from RNA-Seq: A case study of Polytomella non-photosynthetic green algae. Mol Phylogenet Evol. 2016;98:57–62. doi:10.1016/j.ympev.2016.01.017.

73. Waits DS, Simpson DY, Sparkman AM, Bronikowski AM and Schwartz TS. The utility of reptile blood transcriptomes in molecular ecology. Molecular Ecology Resources. 2020;20 1:308–17. doi:10.1111/1755-0998.13110.

74. Kumazawa Y. Mitochondrial DNA sequences of five squamates: phylogenetic affiliation of snakes. DNA Research. 2004;11 2:137–44.

75. Li H and Durbin R. Fast and accurate short read alignment with Burrows-Wheeler transform. Bioinformatics. 2009;25 doi:10.1093/bioinformatics/btp324.

76. Li H, Handsaker B, Wysoker A, Fennell T, Ruan J, Homer N, et al. The Sequence Alignment/Map format and SAMtools. Bioinformatics. 2009;25 16:2078–9. doi:10.1093/bioinformatics/btp352.

77. Katoh K and Standley DM. A simple method to control over-alignment in the MAFFT multiple sequence alignment program. Bioinformatics. 2016;32 13:1933–42. doi:10.1093/bioinformatics/btw108.

78. Kearse M, Moir R, Wilson A, Stones-Havas S, Cheung M, Sturrock S, et al. Geneious Basic: An integrated and extendable desktop software platform for the organization and analysis of sequence data. Bioinformatics. 2012;28 12:1647–9.

79. Pertea M, Kim D, Pertea GM, Leek JT and Salzberg SL. Transcript-level expression analysis of RNA-seq experiments with HISAT, StringTie and Ballgown. Nat Protoc. 2016;11 9:1650–67. doi:10.1038/nprot.2016.095.

80. Picard Toolkit. http://picard.sourceforge.net/. 2019.

81. Leache AD, Harris RB, Maliska ME and Linkem CW. Comparative species divergence across eight triplets of spiny lizards (Sceloporus) using genomic sequence data. Genome Biol Evol. 2013;5 12:2410–9. doi:10.1093/gbe/evt186.

82. Arthofer W, Banbury BL, Carneiro M, Cicconardi F, Duda Thomas F, Harris RB, et al. Genomic Resources Notes Accepted 1 August 2014–30 September 2014. Molecular Ecology Resources. 2014;15 1:228–9. doi:10.1111/1755-0998.12340.

83. Li H. Aligning sequence reads, clone sequences and assembly contigs with BWA-MEM. arXiv. 2013;00 00:1–3. doi:arXiv:1303.3997 [q-bio.GN].

84. McKenna A, Hanna M, Banks E, Sivachenko A, Cibulskis K, Kernytsky A, et al. The Genome Analysis Toolkit: a MapReduce framework for analyzing next-generation DNA sequencing. Genome Research. 2010;20:1297–303.

85. Depristo MA, Banks E, Poplin R, Garimella KV, Maguire JR, Hartl C, et al. A framework for variation discovery and genotyping using next-generation DNA sequencing data. Nature Genetics. 2011;43 5:491–501. doi:10.1038/ng.806.

86. Van der Auwera GA, Carneiro MO, Hartl C, Poplin R, Del Angel G, Levy-Moonshine A, et al. From FastQ data to high confidence variant calls: the Genome Analysis Toolkit best practices pipeline. Curr Protoc Bioinformatics. 2013;43:11 0 1–33. doi:10.1002/0471250953.bi1110s43.

87. Quinlan AR and Hall IM. BEDTools: A flexible suite of utilities for comparing genomic features. Bioinformatics. 2010;26 6:841–2. doi:10.1093/bioinformatics/btq033.

88. Bedoya AM and Leaché AD. Characterization of a large pericentric inversion in Plateau Fence Lizards, (Sceloporus tristichus): evidence from chromosome-scale genomes. bioRxiv. 2020; doi:10.1101/2020.03.18.997676.

89. Castoe TA, de Koning APJ, Hall KT, Card DC, Schield DR, Fujita MK, et al. The Burmese python genome reveals the molecular basis for extreme adaptation in snakes. Proceedings of the National Academy of Sciences. 2013;110 51:20645–50.

90. Rasys AM, Park S, Ball RE, Alcala AJ, Lauderdale JD and Menke DB. CRISPR-Cas9 Gene Editing in Lizards through Microinjection of Unfertilized Oocytes. Cell Rep. 2019;28 9:2288–92 e3. doi:10.1016/j.celrep.2019.07.089.

91. Cox RM, Skelly SL and John-Alder HB. Testosterone Inhibits Growth in Juvenile Male Eastern Fence Lizards (Sceloporus undulatus): Implications for Energy Allocation and Sexual Size Dimorphism. Physiological and Biochemical Zoology. 2005;78 4:531–45.

92. Cox RM and John-Alder HB. Testosterone has opposite effects on male growth in lizards (Sceloporus spp.) with opposite patterns of sexual size dimorphism. J Exp Biol. 2005;208 Pt 24:4679–87. doi:10.1242/jeb.01948.

93. John-Alder HB, Cox RM and Taylor EN. Proximate developmental mediators of sexual dimorphism in size: case studies from squamate reptiles. Integr Comp Biol. 2007;47 2:258–71. doi:10.1093/icb/icm010.

94. Schrey AW, Robbins TR, Lee J, Dukes DW, Ragsdale AK, Thawley CJ, et al. Epigenetic response to environmental change: DNA methylation varies with invasion status. Environmental Epigenetics. 2016;2 2:dvw008. doi:10.1093/eep/dvw008.

